# Low mutation rate but high male-bias in the germline of a short-lived opossum

**DOI:** 10.1101/2024.12.05.627076

**Authors:** Yadira Peña-Garcia, Richard J. Wang, Muthuswamy Raveendran, R. Alan Harris, Paul B. Samollow, Jeffrey Rogers, Matthew W. Hahn

## Abstract

Age and sex have been found to be important determinants of the mutation rate per generation in mammals, but the mechanisms underlying these factors are still unclear. One approach to distinguishing between alternative mechanisms is to study species that reproduce at very young ages, as competing hypotheses make different predictions about patterns of mutation in these organisms. Here, we study the germline mutation rate in the gray short-tailed opossum, Monodelphis domestica, a laboratory model species that becomes reproductively mature at less than six months of age. Whole-genome sequencing of 22 trios reveals the lowest mutation rate per generation found in mammals thus far (0.26 × 10-8 per base pair per generation at an average parental age of 313 days), which is expected given their early reproduction. We also examine the mutation spectrum and find fewer mutations at CpG sites in opossums than in humans, consistent with the lower CpG content in the opossum genome. We observe that two-thirds of mutations are inherited from the male parent in opossums, slightly lower than the degree of male bias observed in organisms that reproduce at much older ages. Nevertheless, the very young age at reproduction in opossums suggests that ongoing spermatogonial divisions in males after puberty are not the primary driver of the observed male mutation bias. These findings contribute to a growing body of evidence that the differences between male and female germline mutation may arise from mechanisms other than cell division post-puberty.

**Article Summary:** This study investigates the germline mutation rate in the gray short-tailed opossum, a marsupial with early reproductive maturity. By sequencing 22 families, we report the lowest mutation rate recorded in mammals but a typical male mutation bias. These findings add to growing evidence that challenges the traditional view that continuing cell division is the primary driver of male-biased mutations. Instead, the study suggests that alternative mechanisms, such as differences in DNA repair, may influence sex-specific mutation rates.

## Introduction

Germline mutations are the primary source of genetic variation and a fundamental driver of evolution. While most *de novo* mutations (DNMs) are neutral, some can cause heritable diseases or contribute to adaptive traits, highlighting their critical role in both evolutionary processes and human health. The first estimates of the mutation rate were derived from the study of rare Mendelian diseases in humans (Haldane 1935) and then later through the analysis of sequence divergence between species with known divergence times (Nachman And Crowell 2000; Silva And Kondrashov 2002). Advances in sequencing technologies have facilitated the direct estimation of mutation rates using pedigrees, enabling the identification of mutations arising between parents and offspring (Yoder And Tiley 2021). As pedigree sequencing proliferates, the number of species for which the mutation rate has been estimated continues to grow (e.g. Venn *et al*. (2014); Thomas *et al*. (2018); Besenbacher *et al*. (2019); Koch *et al*. (2019); Wang *et al*. (2020); Wu *et al*. (2020); Wang *et al*. (2022a); Wang *et al*. (2022b); Bergeron *et al*. (2023); Suarez-Menendez *et al*. (2023); Armstrong *et al*. (2024)). These studies have shown that per-generation mutation rates can vary by a factor of 40 across vertebrates (Bergeron *et al*. 2023).

Variation in the per-generation mutation rate has been attributed to differences in life-history traits, especially generation time (Thomas *et al*. 2018; Wang *et al*. 2022b; Bergeron *et al*. 2023). In mammals, generation time has a large effect because both male and female parents transmit more mutations as they age (Kong *et al*. 2012; Goldmann *et al*. 2016; Wong *et al*. 2016; Jonsson *et al*. 2017). This means that species with shorter generation times will have lower per-generation mutation rates, simply because they reproduce at younger ages. In addition, mammalian fathers consistently pass on more mutations than mothers—approximately three-fourths of all mutations (De Manuel *et al*. 2022)—a phenomenon known as male mutation bias (WILSON SAYRES AND MAKOVA 2011). This difference between the sexes has been attributed to the higher number of cell divisions, and thus DNA replication cycles, during spermatogenesis compared to oogenesis, leading to a greater incidence of replication errors in males; it is thought that the male and female germlines undergo a similar number of cell divisions before puberty (Drost And Lee 1995; Crow 2000). However, studies in humans, mice, and cats have shown the existence of a strong male bias even shortly after puberty, which is not consistent with a replication- driven hypothesis for male bias (Jonsson *et al*. 2017; Lindsay *et al*. 2019; Wang *et al*. 2022b). Instead, alternative hypotheses for the origin of male bias have been proposed (Gao *et al*. 2019; Hahn *et al*. 2023), though more species must be studied in order to properly evaluate alternative models.

Here, we contribute to our understanding of mammalian germline mutation rates and male mutation bias. We used pedigree sequencing data from the gray short-tailed opossum (*Monodelphis domestica*), originally native to Brazil and surrounding countries (Vandeberg AND ROBINSON 1997), to explore the effect of paternal and maternal age on the mutation rate. Marsupials and eutherian mammals diverged approximately 160 million years ago (Luo *et al*. 2011), with marsupials exhibiting many distinct developmental and gestational traits compared to eutherians. While there are multiple studies of the effects of parental age and male-biased mutation among eutherians, we present the first detailed account in a marsupial. These data therefore provide a unique test of how well mutation models developed within Eutheria fit across mammals. Our results in the opossum reveal the lowest mutation rate per site per generation reported to date in mammals, aligning with expectations given the early age of reproduction among samples in our study (0.5-1 year). We also observe a significant male mutation bias, even among the youngest parents.

Finally, our analysis of the mutation spectrum showed a difference from other mammals, especially in mutations at CpG dinucleotides. These findings provide new insights into the cellular mechanisms governing the evolution of mutation rates and sex-specific biases in mammals, and emphasize the importance of studying diverse species to understand the broad applicability of existing models.

## Materials and Methods

### Samples and sequencing

Liver samples were collected from a total of 34 individuals belonging to three extended pedigrees, comprising 22 trios, of gray short-tailed opossums (*Monodelphis domestica*) kept in captivity at Texas A&M University (College Station, TX). Animal care and experimental procedures were conducted in strict accordance with policies and guidelines of Texas A&M. Genomic DNA extraction was performed from the collected samples using Gentra Puregene methods and reagents, following established protocols. The extracted DNA was sheared into fragments of approximately 200-600 bp. Sequencing libraries were prepared as described in Wang *et al*. (2022b). Briefly, PCR-free libraries were generated using KAPA Hyper PCR-free reagents. The sheared DNA molecules were then purified using AMPure XP beads. The procedure next included DNA end-repair and 3’ adenylation followed by addition of barcoded adapters, generating paired-end reads with an average length of 426 bp. Whole genome sequencing of the prepared libraries was performed on Illumina Hi-Seq X instruments, following manufacturer’s recommendation with minor modifications. This sequencing approach provided ample coverage of the opossum genome (average 35x), enabling accurate detection and characterization of germline mutations within the pedigrees.

### Read mapping and variant calling

Sequencing reads obtained from all 34 samples were mapped to the opossum reference genome MonDom5 (NCBI RefSeq GCF_000002295.2; (Mikkelsen *et al*. 2007)) using BWA-MEM v. 0.7.12-r1039 (Li 2013). Duplicate reads were identified and marked using Picard MarkDuplicates (v2.6.0; http://broadinstitute.github.io/picard/) to ensure accurate variant calling. Single nucleotide variants (SNVs) were called using the Genome Analysis Toolkit (GATK) version 4.1.2.0 (Van Der Auwera *et al*. 2013) following recommended best practices. HaplotypeCaller was employed to generate genomic Variant Call Format (gVCF) files for each individual sample. This approach allows for the identification of variant sites along with genotype likelihoods. Subsequently, joint genotype calling across all samples was performed with GenotypeGVCFs, facilitating the comprehensive assessment of genetic variation within the opossum population. To ensure the integrity of the variant dataset, we applied stringent filtering criteria. Specifically, we implemented filters recommended by GATK, targeting single nucleotide polymorphisms (SNVs) with parameters set as follows: "QD < 2.0 || MQ < 40.0 || FS > 60.0 || SOR > 3.0 || MQRankSum < -12.5 || ReadPosRankSum < -8.0 and removed calls that failed.

### Identification of candidate mutations

We followed a framework outlined in our prior studies (Wang *et al*. 2020; Wang *et al*. 2022a; Wang *et al*. 2022b) to detect autosomal *de novo* mutations from the set of called variants. Briefly, we restricted our search to a subset of sites identified as “Mendelian violations” within each trio by GATK. These are sites where both parents are homozygous for the reference allele, while the offspring is heterozygous for an alternate allele. To ensure the selection of high-confidence candidate mutation, we implemented a series of stringent filtering criteria:

- Read-depth filtering: We restricted candidate sites to those with a read-depth ranging between 20 and 60 for each individual within the trio. This step aims to mitigate sampling errors associated with inadequate coverage and addresses potential issues resulting from repetitive genomic regions.
- Genotype quality assessment: All individuals were subjected to stringent genotype quality thresholds (GQ > 70), ensuring reliable genotype calls across the cohort.
- Strand-specificity check: We required that candidate mutations were observed on both the forward and reverse strands in the offspring, as indicated by allelic depth metrics (ADF, ADR > 0).
- Parental validation: Mutations with more than one alternate read on either strand in nominally homozygous parents (AD = 0) were excluded to minimize genotyping errors and to ensure that selected variants were not inherited from parental sources (i.e. the parent is actually heterozygous). The original BAM files were checked for such reads.
- Pedigree exclusivity: Candidate mutations should exclusively be observed in samples directly descended from the parental lineage within each trio and not in any other individual in the population.
- Allelic balance filtering: Candidate mutations were further refined based on allelic balance criteria (Allelic Balance > 0.3), ensuring a balanced representation of alternate and reference alleles in the offspring.

Our aim in implementing stringent filters was to minimize the occurrence of false-positive discoveries while ensuring the robustness of our mutation rate estimate.

### Estimating the per-generation mutation rate

To estimate the per-generation mutation rate accurately, the counts of *de novo* mutations identified are divided by the total number of callable sites in each trio. Thus, we first defined the callable genome, the subset of the entire genome where reliable variant calls can be made with high confidence. We applied a methodology similar to the one used in previous studies (Besenbacher *et al*. 2019; Wang *et al*. 2020; Wang *et al*. 2022a; Wang *et al*. 2022b). In these studies, a callable site is defined as one where sequencing depth and quality meet predefined criteria, so that the estimated probability of an identified candidate mutation is a true mutation is high. The mutation rate calculation follows the equation:

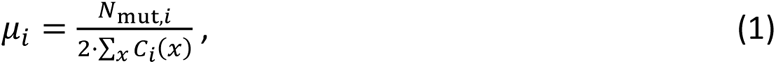

where 𝜇_𝑖_ is the per-site per generation mutation rate for trio 𝑖, 𝑁_mut,𝑖_is the number of *de novo* mutations identified in trio 𝑖, and 𝐶_𝑖_(𝑥) is the callability of site 𝑥 in the trio. This method assumes the independence of calling each individual in the trio correctly, which enables us to estimate 𝐶_𝑖_(𝑥) as the product of the probabilities of calling the child, father, and mother correctly as 𝐶_𝑐_(𝑥), 𝐶_𝑝_(𝑥), and 𝐶_𝑚_(𝑥), respectively, so that:

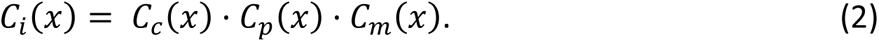

These probabilities are determined by applying the same set of stringent filters to high-confidence heterozygous calls from each trio. For heterozygous SNVs in the offspring, the callability 𝐶_𝑐_(𝑥) was estimated as the ratio of filtered heterozygous variants, 𝑁_het,filtered_, to all heterozygous variants, 𝑁_het,all_, where 𝑁_het,all_, represents the number of variants in the offspring where one parent is homozygous for the reference, and the other parent is homozygous for the alternate allele. These numbers lead to an estimate of callability in the child:

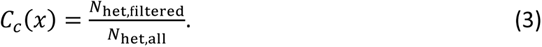

Similarly, parental callability, 𝐶_𝑝_(𝑥) and 𝐶_𝑚_(𝑥), was estimated by determining the proportion of remaining sites after applying stringent mutation filters. As implemented by (Wang *et al*. 2020; 2022a; 2022b), we evaluated the callability for each individual using a random sample of 250,000 sites across the genome that matched the respective criteria for each trio.

### Prediction of mutation rate from human data

To assess whether the observed mutation rates in opossums were consistent with expectations based on parental age, we utilized parameters estimated from human data (Jonsson *et al*. 2017). This model estimates the maternal and paternal contribution to the overall mutation rate in a trio, based on the age at conception of the parents.

Assuming a constant accumulation of mutations across the lifespan of the parents, we model the paternal and maternal contributions to a single offspring as follows:

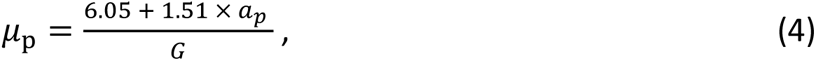

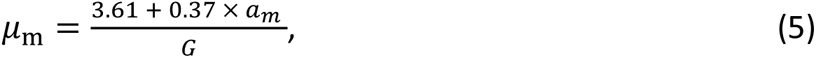

where 𝑎_𝑝_ and 𝑎_𝑚_represent the paternal and maternal ages at conception (in years), respectively, and *G* is the callable genome size used by Jonsson et al. (=2.68 × 10⁹ bp). The total predicted per-generation mutation rate is then (𝜇_p_ + 𝜇_m_)/2 for a set of specified parental ages.

### Assigning mutations to a parent of origin

To determine the parental origin of the *de novo* mutations identified in the 22 opossum trios, we utilized two tools: POOHA (https://github.com/besenbacher/POOHA; (Maretty *et al*. 2017; Besenbacher *et al*. 2019; Bergeron *et al*. 2021); ) and Unfazed (Belyeu *et al*. 2021).

POOHA assigns the parent of origin by analyzing informative heterozygous single nucleotide polymorphisms (SNPs) that are located within the same sequencing read or read-pair as the DNM. We ran POOHA with a 1000 bp window around each DNM, using the variant call format (VCF) file for each trio and a BAM file for the child only.

In contrast, Unfazed implements a broader strategy, utilizing SNPs beyond the immediate vicinity of the DNM by linking them to parent-specific markers through haplotype reconstruction. Unfazed can extend the region of informative SNPs by chaining reads together using mutually overlapping heterozygous sites, allowing for assignment over longer distances than single read-pairs typically permit. This makes Unfazed particularly effective in cases where the parental source of the mutation is not defined directly by a nearby SNP. For each DNM, we provided Unfazed with the same 1000 bp window around each site as was used for POOHA, and the software used this window to search for informative sites.

To ensure accuracy, we cross-referenced the results from both tools, manually reviewing all assigned mutations using the Integrative Genomics Viewer (IGV) (Robinson *et al*. 2023).

## Results

### Estimating the mutation rate from opossum pedigrees

The gray short-tailed opossum becomes sexually mature at 5-6 months and can live for 36- 42 months in captivity (Macrini 2004). All individuals in this study reproduced before they were 17 months old (Table 1). In total, we sequenced the genomes of 34 individuals from 3, three-generation extended pedigrees (Figure 1) of captive opossums maintained at Texas A&M University. Genomic DNA was isolated and sequenced on Illumina Hi-Seq X instruments, producing 150 bp paired-end reads to an average of 35× coverage (min: 31×, max: 38×) (Table 1; Materials and Methods). The pedigrees can be separated into 22 trios, and we estimated the mutation rate for each trio independently. Filtering strategies were used to retain only sites with optimal quality and coverage, which resulted in an average callable genome of 2.2 Gb per trio (Table 1).

**Figure 1.**
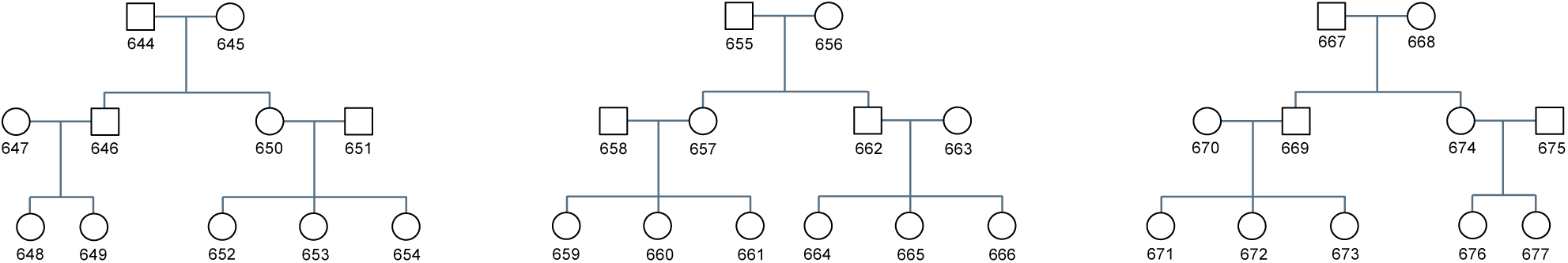
Pedigree of sequenced gray short-tailed opossum individuals. Three extended pedigrees consisting of 34 individuals total were collected, which can be divided into 22 trios. Males are represented by squares and females by circles. IDs of all individuals correspond to those listed in Table 1.

**Table 1.** Summary of mutation counts and mutation rate per trio.

| Offspring ID | Parental age at conception (days) | | Mutations | Phased mutations | Mutations of paternal origin | Mutations of maternal origin | Mean read depth | Callable genome size (Mb) | Mutation rate ($\times 10^{-8}$ /bp) |
| --- | --- | --- | --- | --- | --- | --- | --- | --- | --- |
|  | Paternal | Maternal |  |  |  |  |  |  |  |
| 646 | 286 | 167 | 7 | 7 | 4 | 3 | 35.3 | 2201.29 | 0.159 |
| 648 | 190 | 167 | 16 | 11 | 10 | 1 | 35.0 | 2153.665 | 0.371 |
| 649 | 270 | 247 | 9 | 8 | 5 | 3 | 36.0 | 2220.75 | 0.203 |
| 652 | 298 | 345 | 6 | 4 | 2 | 2 | 36.0 | 2215.335 | 0.135 |
| 653 | 398 | 445 | 13 | 8 | 5 | 3 | 36.3 | 2256.59 | 0.288 |
| 654 | 484 | 531 | 20 | 17 | 11 | 6 | 36.0 | 2071.365 | 0.483 |
| 657 | 393 | 252 | 7 | 7 | 2 | 5 | 32.7 | 2059.23 | 0.17 |
| 659 | 168 | 180 | 14 | 12 | 6 | 6 | 35.0 | 2222.33 | 0.315 |
| 660 | 293 | 305 | 15 | 10 | 7 | 3 | 35.0 | 2275.71 | 0.33 |
| 661 | 372 | 384 | 7 | 5 | 3 | 2 | 34.7 | 2225.295 | 0.157 |
| 664 | 237 | 197 | 10 | 7 | 7 | 0 | 34.3 | 2178.465 | 0.23 |
| 665 | 362 | 322 | 15 | 9 | 3 | 6 | 34.7 | 2264.995 | 0.331 |
| 666 | 472 | 432 | 12 | 9 | 7 | 2 | 34.7 | 2216.325 | 0.271 |
| 669 | 328 | 146 | 9 | 9 | 8 | 1 | 34.7 | 2206.54 | 0.204 |
| 671 | 278 | 209 | 10 | 7 | 7 | 0 | 34.7 | 2200.54 | 0.227 |
| 672 | 390 | 321 | 10 | 4 | 3 | 1 | 34.3 | 2198.17 | 0.227 |
| 673 | 504 | 435 | 10 | 8 | 4 | 4 | 34.3 | 2174.16 | 0.23 |
| 676 | 198 | 293 | 12 | 10 | 5 | 5 | 36.3 | 2200.665 | 0.273 |
| 677 | 323 | 418 | 20 | 15 | 12 | 3 | 35.7 | 2167.075 | 0.461 |
| 650 | 286 | 167 | 7 | 6 | 4 | 2 | 35.3 | 2186.755 | 0.16 |
| 662 | 393 | 252 | 7 | 6 | 4 | 2 | 33.0 | 2071.105 | 0.169 |
| 674 | N/A | N/A | 10 | 9 | 6 | 3 | 35.0 | 2153.915 | 0.232 |

After stringent filtering to reduce the incidence of false positives, we identified a total of 246 single nucleotide *de novo* mutations across the 22 trios, four of which were multinucleotide mutations (Schrider *et al*. 2011). The number of DNMs identified in each trio ranged from as low as 6 to as high as 20 (Table 1). None of the mutations were recurrent across unrelated individuals, but 12 mutations (6 genomic positions) were shared between 5 pairs of siblings, implying that these mutations were mosaic in the germline of a parent. We also tracked the transmission of DNMs across generations within our pedigrees for the six individuals in the first generation that went on to become parents. A total of 47 detected mutations could be tracked, of which 43.6%, on average, were transmitted to offspring in the next generation (Table S1); this transmission rate is not different from the 50% expectation (*P*=0.18, Exact test). These results help to strengthen the inference that these are true DNMs.

Considering all trios and DNMs, we estimated an average mutation rate of 0.256 × 10^-8^ per base pair per generation (95% confidence interval [CI]: 0.214–0.298 × 10^-8^) for parents at an average age of 313 days across sexes (♂:330 days ♀:296 days). In humans, per-generation estimates are 1.29 × 10^-8^ per base pair with an average parental age of 30.1 years (Jonsson *et al*. 2017). Using regression estimates based on human data (Methods) and applying them to opossums after adjusting for differences in the sizes of the callable genomes, we predict an average mutation rate of 0.211 × 10^-8^ per base pair per generation (95% CI: 0.207–0.215 × 10^-8^) for humans that have children at the same age as the average opossum. This prediction closely matches our observed mutation rate (and is within its CI) (Figure S1), further highlighting the significant role of parental age in determining mutation rates across different species.

### Parental contributions to the mutation rate

We used two tools, POOHA (Maretty *et al*. 2017; Bergeron *et al*. 2021) and Unfazed (Belyeu *et al*. 2021), to determine the parent of origin for DNMs across the 22 trios (sometimes this process is called “phasing”). POOHA assigned 193 of the 246 mutations, while Unfazed assigned only 170 mutations. Although Unfazed also successfully assigned two mutations that POOHA could not, both programs assigned the same parent of origin for all shared mutations, with no disagreements. To ensure that all the mutations were assigned correctly, we manually inspected all assigned mutations in IGV. We found two errors from Unfazed, whereas POOHA assigned five mutations incorrectly. Therefore, we excluded these cases from the set of mutations with determined parental origin in our subsequent analyses. However, despite these minor discrepancies, both tools provided reliable results overall. These steps resulted in 188 mutations (76.4%) assigned by POOHA, and 168 mutations (68.4%) assigned by Unfazed. Based on the larger number of mutations assigned to one parent or the other, we proceeded with the POOHA results for the remainder of our analyses.

Of the 188 assigned mutations, 125 (66.5%) were of paternal origin and 63 (33.5%) were of maternal origin (Table 1). This indicates a strong male mutation bias, with nearly two-thirds of mutations being paternal. However, this bias is slightly lower than the 80.4% reported in humans (Jonsson *et al*. 2017) and the 75-80% observed in other mammals (Tatsumoto *et al*. 2017; Wang *et al*. 2022a; Wang *et al*. 2022b). We find that our observed fraction of paternal mutations is significantly lower than that among humans (c^2^=21.80; *P*<0.00001) and significantly lower than an expectation of 75% (c^2^=7.26; *P*=0.007).

In contrast to what has been observed in other mammals (Venn *et al*. 2014; Rahbari *et al*. 2016; Thomas *et al*. 2018; Sasani *et al*. 2019; Wang *et al*. 2020; Wang *et al*. 2022b), we did not find a parental age effect on the assigned mutations (Figure 2). Neither paternal age nor maternal age had a significant effect on either the total number of mutations or the number of sex-specific mutations (i.e. an effect of maternal age on maternally transmitted mutations).

**Figure 2.**
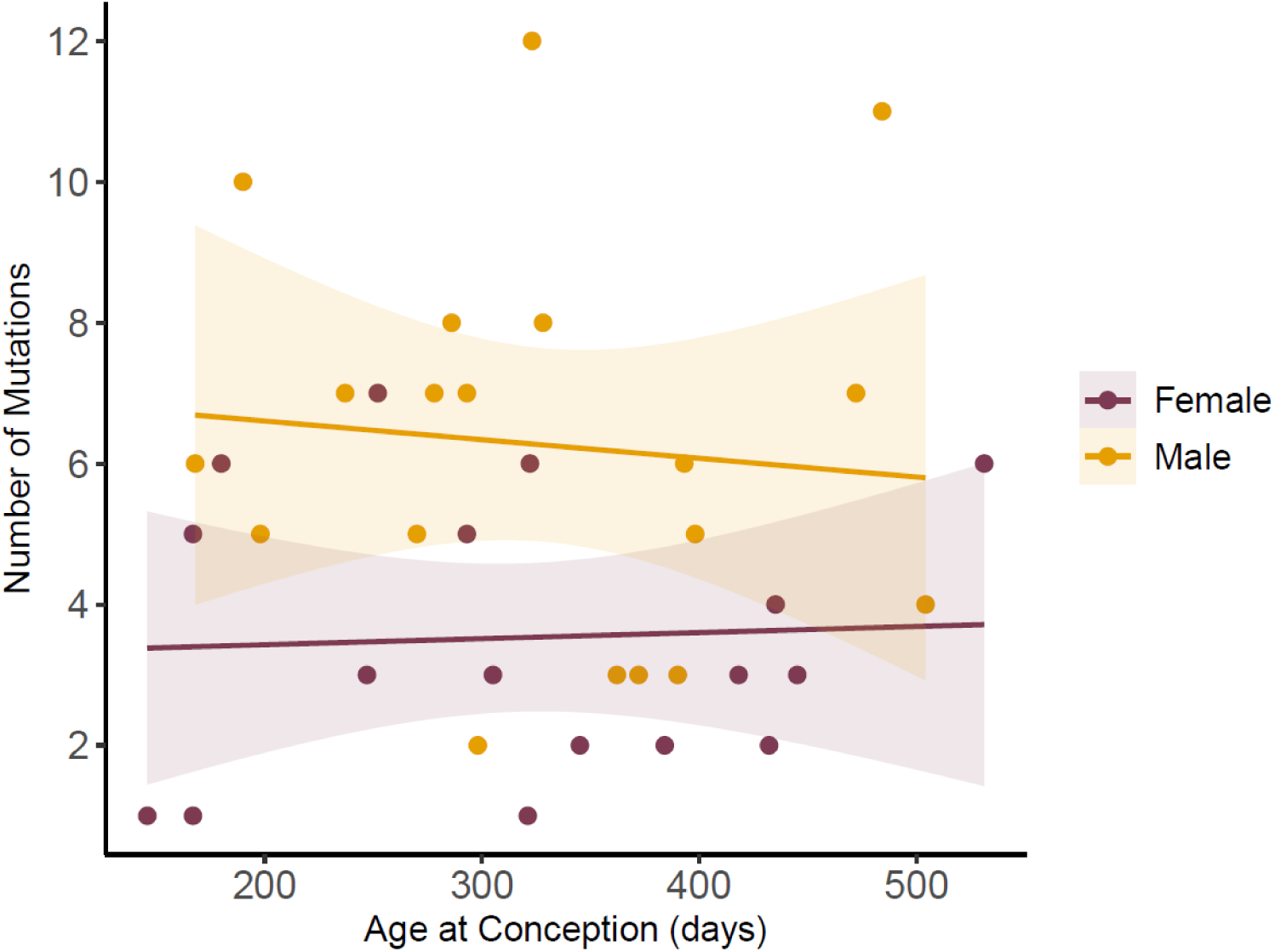
Number of mutations as a function of sex and parental age. Each dot represents the number of mutations found in a single offspring, color-coded by parent of origin: yellow for mutations from the male parent and maroon for mutations from the female parent. The solid lines are the best-fit regression for the relationship between parental age and mutation count, with shaded areas representing the 95% confidence intervals.

One potential explanation for the lack of a parental age effect could be the small age-range of parents in our study: the youngest parent differs by only one year from the oldest (Table 1; Figure 2). To determine whether this narrow age range could be the reason why we did not observe a trend in opossums, we evaluated the same age-range restrictions in a human dataset. Using the data from Jonsson *et al*. (2017), we analyzed pairs of datapoints one year apart (the human data only has resolution to the nearest year). We identified 44 such pairs, where consecutive ages were available in the dataset, and then asked whether there was a significant effect of paternal age on the number of paternally transmitted mutations. Only 7 out of 44 pairs showed a significant paternal age effect (Figure S2), even though there is a strong effect when analyzing all the data together (see Figure 1 in Jonsson et al. 2017). These results suggest that an age effect in opossums as strong as the one in humans might not be detected due to the narrow age range among parents in our opossum sample.

### Mutation spectrum in opossum

We examined the mutation spectrum from all identified mutations, classifying each into one of six mutation types (Figure 3A). While the frequency of some mutation classes closely resembled those found in humans (Figure 3B), the overall mutation spectrum was significantly different between the two species (c^2^=30.69; *P*=0.000011). The transition-to-transversion (Ts/Tv) ratio in opossum was 1.83, comparable to the ratio observed in other mammals, and not significantly different than the human ratio of 2.22 (Jonsson *et al*. 2017) (c^2^=1.95;*P=*0.16) C > T transitions were the most common type of mutation, observed in 44% (108 out of 246) of all changes, similar to the 42.1% reported in humans (Jonsson *et al*. 2017).

**Figure 3.**
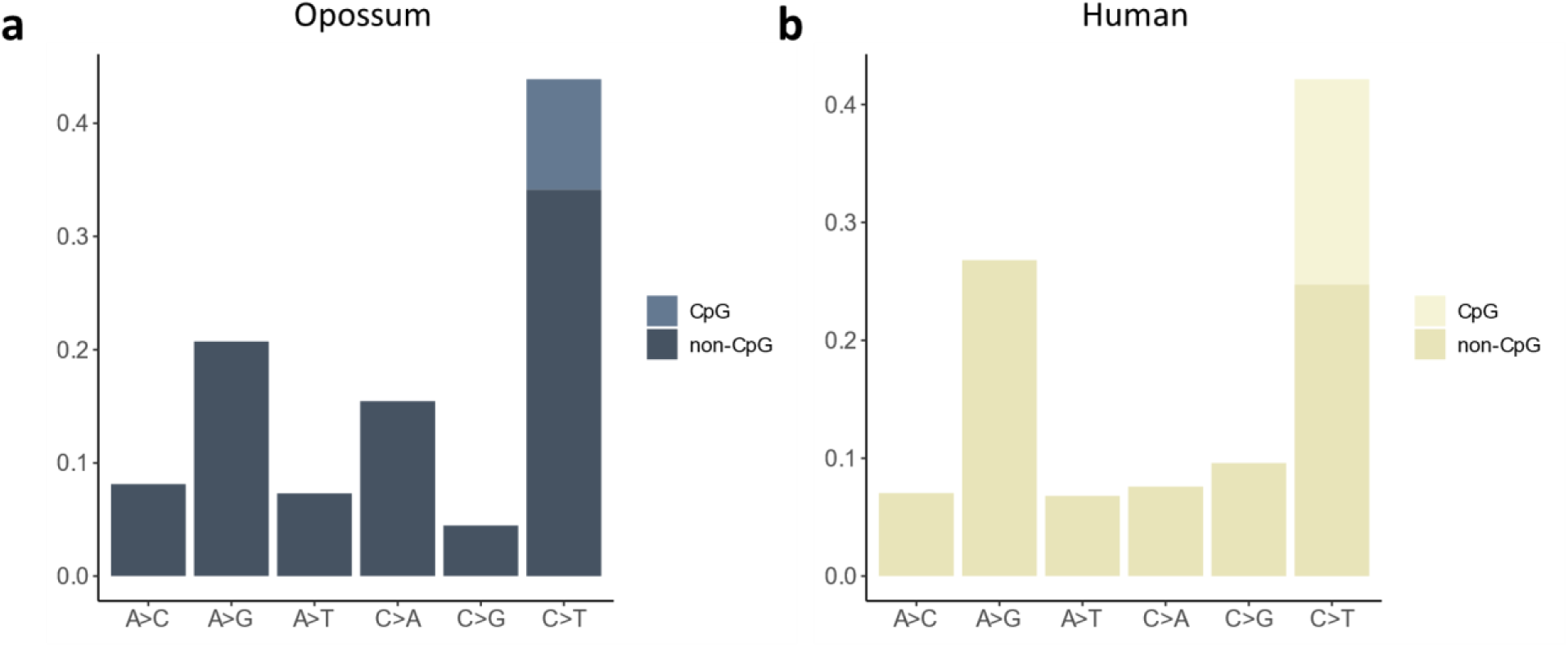
Opossum mutation spectrum compared to humans. **(a)** The proportion of each mutation class observed among gray short-tailed opossum trios, including their reverse complements. Mutation classes are categorized based on nucleotide changes (e.g., C→T, G→A). **(b)** The same mutation spectrum for human trios, highlighting similarities and differences in mutation patterns between the two species. Data on humans from Jonsson *et al*. (2017).

However, only 9.8% of the total number of mutations arose at CpG sites, lower than the 20.8% of all mutations estimated in humans (Besenbacher *et al*. 2015) or 21% in cats (Wang *et al*. 2022b). One possible explanation for the lower fraction of CpG mutations is that the CpG content in the opossum genome is low. At 0.44%, the CpG content in the opossum genome assembly (MonDom5) is less than half of what it is found to be in the human genome (hg38), 1.03%. Though CpG sites are a smaller fraction of the opossum genome, we estimate the mutation rate at these sites to be 5.67 × 10^-8^ per bp per generation, 22.2× higher than the overall rate, and similar to the multiple for CpG sites relative to the overall rate in both humans and cats (21.4× and 21.7×, respectively; (Besenbacher *et al*. 2015; Wang *et al*. 2022b)).

To ask whether the difference in mutation spectrum between humans and opossums is solely due to the lower number of CpG mutations in opossums, we reanalyzed the data excluding CpG sites. Even after removing this group, the mutation spectrum remained significantly different between the two species (c^2^=34.40; *P*= 0.000002). To further explore whether the mutation spectrum observed in opossums could be explained by their age at reproduction, we applied a method previously described in Wang *et al*. (2022b). This approach, originally developed to model the mutation patterns in cats relative to humans, uses Poisson regression to predict the mutation spectrum based on parental age. For each mutation class, a regression model estimates the counts as a function of the father’s and mother’s age at conception, accounting for the fact that mutation rates typically follow a Poisson distribution. When applied to our opossum mutation data, the predicted mutation spectrum still differed substantially from the one we observed (c^2^=11.82; *P*=0.037) (Figure S3). However, this difference was smaller compared to the observed human mutation spectrum, suggesting that while some aspects of the mutation spectrum have evolved between humans and opossums, accounting for dinucleotide content and parental age at conception reduces the discrepancy.

To evaluate variation in the types of mutations inherited from mothers versus fathers, we analyzed the sex-specific mutation spectrum (Figure S4). We observed that the Ts/Tv ratio for males was 2.29, while for females it was lower at 1.42. Similarly, C>T transitions, the most common mutation type, accounted for 48.0% of paternal mutations and 34.9% of maternal mutations. The occurrence of C>T transitions at CpG sites was also less frequent in maternal mutations, at 4.8%, compared to 10.4% in paternal mutations. While none of these differences reached statistical significance (*p* > 0.05), they suggest potential sex-specific trends in the mutation spectrum that merit further exploration.

## Discussion

The results presented here provide valuable insights into the evolution of mutation rates and sex-biased mutation. We identified a total of 246 germline mutations in 22 opossum trios, which enabled us to estimate an average mutation rate of 0.256 × 10^-8^ per base pair per generation for parents at an average age of 313 days. This estimate of the mutation rate in opossums is the lowest value observed so far in mammals (Bergeron *et al*. 2023). A previous estimate derived from a single trio of the same species had a value of 0.46 × 10^-8^ (Bergeron *et al*. 2023), but the parents in that trio had an average age of 1.5 years, which could explain the higher rate observed.

The age at conception appears to be the primary determinant of both the mutation rate per generation (Thomas *et al*. 2018; Bergeron *et al*. 2023) and the mutation spectrum (Wang *et al*. 2022b; Beichman *et al*. 2023) among mammals. These relationships are reasonable when we consider that both the number of mutations and the spectrum of mutations change with the age of parents within species (Jonsson *et al*. 2017). Given the association between age at conception and mutation rate, one can ask whether the rate we observe in opossums is consistent with expectations. Using Poisson regression estimates for the number of mutations expected from male and female parents in humans (Jonsson *et al*. 2017), we predicted the mutation rate in opossums based on the observed parental ages in our trios. The model, adjusted for differences between the callable genome sizes in humans and opossums, predicted a mutation rate of 0.211 × 10⁻^8^ per base pair per generation. This estimate closely matches the observed mutation rate of 0.256 × 10⁻^8^, with no statistically significant difference based on a comparison with the 95% CI of the estimated rate.

Although opossums show the lowest mutation rate among mammals so far, they are not the species with the youngest parents tested: laboratory mice are (Lindsay *et al*. 2019; Lopez-Cortegano *et al*. 2024). Previous work on commonly used inbred strains have reported mutation rates ranging from 0.39 × 10⁻⁸ per base pair per generation for individuals with an average parental age of 171.5 days (Lindsay *et al*. 2019) to 0.67 × 10⁻⁸ per base pair per generation for parents with an average age of approximately 150 days (Lopez-Cortegano *et al*. 2024).These estimates are higher than predicted by our model given the age of the parents, making laboratory mice an outlier. The higher mutation rates may possibly be due to the effects of inbreeding in this model organism, a fact that should be considered when including lab-adapted organisms in comparative studies.

Our analysis of mutations transmitted by each parent revealed a marked male mutation bias, even among very young parents, with fathers passing on two-thirds of the mutations observed in the offspring. Pedigree-based studies, reliant on observing transmitted germline mutations among individuals that have reproduced, face limitations in assessing potential sex biases before puberty. Consequently, our understanding of male bias is limited by the age of reproductive maturity at which individuals can contribute genetic material to the next generation. The gray short-tailed opossum, with its early onset of puberty compared to many mammals, provides an ideal organism to study male bias.

Replication errors have long been thought to be a primary source of germline mutations (Haldane 1947), especially as the male germline undergoes many more replications post-puberty. If replication were the primary driver, one might expect a much weaker bias among younger fathers, as males and females will have undergone a similar number of replications before producing their gametes. Although we do observe a slightly reduced male bias in opossums—most mammals inherit three-quarters of mutations through the male parent (De Manuel *et al*. 2022)—the reduced male bias observed in opossums may reflect a combination of biological, evolutionary, and ecological factors. The early reproductive onset in opossums likely limits the time available for differential mutation accumulation between males and females. If replication is a major driver of male bias, male opossums may simply not have had as much time to accumulate mutations relative to females. This timing could reduce the extent of male bias that might otherwise become more evident with age, as seen in other mammals.

Additionally, the evolutionary divergence between marsupials and eutherian mammals may have resulted in distinct mechanisms underlying differences between the sexes. The long evolutionary separation between these lineages could have led to unique changes in gametogenesis or germline maintenance among marsupials, producing sex-specific mutation patterns distinct from those of eutherian mammals. Environmental factors, such as differences in mutagen exposure between sexes, may also shape the observed bias.

Male opossums, like other mammals, may experience greater environmental stresses due to behavioral or ecological roles that increase contact with mutagens. However, in a species with shorter generational cycles, the impact of such exposure may be constrained by time.

Despite the reduced male bias in opossums, our results are still quite similar to those in other species with much longer generation times. This study, therefore, contributes to a growing body of evidence that additional factors beyond DNA replication play a substantial role in shaping the observed bias (Gao *et al*. 2019; De Manuel *et al*. 2022), including possible differences in DNA repair mechanisms, environmental exposures, or selective pressures acting on the male and female germlines (Hahn *et al*. 2023). Of course, the slightly depressed degree of male-bias in opossums could be explained by a relative lack of male germline replication given the short time between puberty and reproduction, but it would only then explain a small fraction of this bias.

We found a significantly different overall mutation spectrum in opossums compared to that reported for humans, consistent with lineage-specific changes in mutational machinery. Among the differences, a notable one is observed in the fraction of C>T mutations occurring at CpG sites, which is lower in opossums compared to humans. An important factor contributing to the reduced proportion of C>T mutations at CpG sites in opossums may be their relatively low CpG content compared to other mammalian species. Specifically, opossums harbor less than half the CpG content found in humans (Mikkelsen *et al*. 2007), suggesting that CpG depletion could play a significant role in shaping the mutation spectrum. This highlights the significance of genome composition and context in shaping mutation patterns across species. Thus, while variation in generation time may provide insights into certain aspects of mutational spectra (e.g. Jonsson *et al*. (2017); Wang *et al*. (2023)), the low CpG content in opossums underscores the importance of considering species-specific genomic features when interpreting mutation patterns.

In conclusion, further research into the molecular processes underlying germline mutations, particularly during early stages of reproductive maturity, will be essential for unraveling the intricate interplay of factors shaping mutation patterns in mammals. By using a model short-lived marsupial, our research provides valuable insights into mutation rates, male bias, and evolutionary implications, both within marsupials and across mammals. The findings presented here therefore contribute to a broader understanding of the evolutionary processes underlying key evolutionary traits.

## Data availability

The raw sequencing reads are available from the NCBI SRA database under accession number PRJNA1139788, https://www.ncbi.nlm.nih.gov/bioproject/?term=PRJNA1139788.

## Author contributions

Y.P.G., R.J.W., J.R., and M.W.H. conceived analyses; Y.P.G., R.A.H., and R.J.W., performed analyses; P.B.S. provided the opossum samples; M.R. extracted DNA and made sequencing libraries; M.W.H. and J.R. supervised sequencing and analysis.

## Acknowledgments

We wish to thank Donna Muzny, Harshavardhan Doddapaneni, Richard Gibbs and their teams at the Human Genome Sequencing Center, Baylor College of Medicine for expert genome sequencing in support of this study. John VandeBerg provided helpful comments on a draft of the manuscript.

