## Supplementary material for "Low mutation rate but high male-bias in the germline of a short-lived opossum": Table S1

**Table S1.** Comprehensive list of identified mutations in opossums.

| CHROM | POS | BCM ID | SEX | REF | ALT | Unfazed | POOHA | FATHER AGE AT CONCEPTION | MOTHER AGE AT CONCEPTION |
|---|---|---|---|---|---|---|---|---|---|
| 1 | 22373653 | 100674 | F | A | G | NA | M | NA | NA |
| 1 | 29274266 | 100674 | F | A | G | P | P | NA | NA |
| 1 | 35444119 | 100648 | F | A | T | P | P | 190 | 167 |
| 1 | 36444744 | 100673 | F | G | A | M | M | 504 | 435 |
| 1 | 42352427 | 100660 | F | C | G | P | P | 293 | 305 |
| 1 | 48364989 | 100648 | F | G | A | P | P | 190 | 167 |
| 1 | 66992151 | 100664 | F | G | A | P | P | 237 | 197 |
| 1 | 74574592 | 100672 | F | G | T | M | M | 390 | 321 |
| 1 | 77016929 | 100646 | M | G | A | P | P | 286 | 167 |
| 1 | 79673462 | 100665 | F | T | C | P | P | 362 | 322 |
| 1 | 84875189 | 100673 | F | G | A | P | NA | 504 | 435 |
| 1 | 86710665 | 100671 | F | G | A | P | P | 278 | 209 |
| 1 | 96974586 | 100665 | F | A | C | M | M | 362 | 322 |
| 1 | 115826679 | 100669 | M | G | A | P | P | 328 | 146 |
| 1 | 127013662 | 100648 | F | A | T | NA | NA | 190 | 167 |
| 1 | 130133628 | 100677 | F | C | T | P | P | 323 | 418 |
| 1 | 141892256 | 100653 | F | A | G | NA | NA | 398 | 445 |
| 1 | 151438228 | 100677 | F | G | A | P | P | 323 | 418 |
| 1 | 170571441 | 100648 | F | C | A | NA | NA | 190 | 167 |
| 1 | 181628090 | 100646 | M | T | C | P | P | 286 | 167 |
| 1 | 181628090 | 100650 | F | T | C | P | P | 286 | 167 |
| 1 | 188888172 | 100662 | M | A | T | M | M | 393 | 252 |
| 1 | 204545521 | 100662 | M | A | G | M | M | 393 | 252 |
| 1 | 210265895 | 100650 | F | C | T | M | M | 286 | 167 |
| 1 | 211720848 | 100677 | F | G | A | P | P | 323 | 418 |
| 1 | 212450662 | 100677 | F | G | T | P | P | 323 | 418 |
| 1 | 260931482 | 100673 | F | G | A | P | P | 504 | 435 |
| 1 | 294059773 | 100673 | F | G | T | M | M | 504 | 435 |
| 1 | 327787778 | 100660 | F | C | T | NA | NA | 293 | 305 |
| 1 | 359096681 | 100664 | F | T | C | P | P | 237 | 197 |
| 1 | 367908729 | 100669 | M | C | T | P | P | 328 | 146 |
| 1 | 371924934 | 100654 | F | G | A | P | P | 484 | 531 |
| 1 | 387967624 | 100653 | F | G | T | P | P | 398 | 445 |
| 1 | 395600782 | 100654 | F | T | C | M | M | 484 | 531 |
| 1 | 415023005 | 100664 | F | C | T | P | P | 237 | 197 |
| 1 | 423898045 | 100672 | F | G | A | NA | NA | 390 | 321 |
| 1 | 436693871 | 100660 | F | T | G | NA | NA | 293 | 305 |
| 1 | 474446409 | 100665 | F | G | A | NA | NA | 362 | 322 |

| | | | | | | | | | |
|---|---|---|---|---|---|---|---|---|---|
| 1 | 488409705 | 100661 | F | G | T | M | M | 372 | 384 |
| 1 | 489675670 | 100673 | F | A | G | P | P | 504 | 435 |
| 1 | 494403956 | 100648 | F | G | A | P | P | 190 | 167 |
| 1 | 508995015 | 100659 | F | G | A | NA | NA | 168 | 180 |
| 1 | 518072699 | 100648 | F | A | T | P | P | 190 | 167 |
| 1 | 529229050 | 100674 | F | G | A | NA | NA | NA | NA |
| 1 | 553941109 | 100654 | F | C | T | P | P | 484 | 531 |
| 1 | 569039872 | 100674 | F | A | G | P | P | NA | NA |
| 1 | 571266583 | 100665 | F | G | A | NA | NA | 362 | 322 |
| 1 | 571266583 | 100666 | F | G | A | P | P | 472 | 432 |
| 1 | 589302113 | 100653 | F | G | C | P | P | 398 | 445 |
| 1 | 597108505 | 100650 | F | C | A | P | P | 286 | 167 |
| 1 | 614721917 | 100660 | F | G | A | M | M | 293 | 305 |
| 1 | 631360174 | 100669 | M | T | G | P | P | 328 | 146 |
| 1 | 639996303 | 100672 | F | G | T | NA | NA | 390 | 321 |
| 1 | 639996303 | 100673 | F | G | T | M | M | 504 | 435 |
| 1 | 662140185 | 100648 | F | C | A | M | M | 190 | 167 |
| 1 | 668601161 | 100672 | F | C | T | P | P | 390 | 321 |
| 1 | 672009526 | 100649 | F | G | A | NA | NA | 270 | 247 |
| 1 | 679326287 | 100652 | F | T | C | P | P | 298 | 345 |
| 1 | 693597222 | 100662 | M | G | C | P | P | 393 | 252 |
| 1 | 734804797 | 100648 | F | C | A | NA | P | 190 | 167 |
| 1 | 736134992 | 100654 | F | C | T | P | P | 484 | 531 |
| 2 | 107867 | 100657 | F | G | A | M | M | 393 | 252 |
| 2 | 4186660 | 100661 | F | C | T | P | P | 372 | 384 |
| 2 | 10207680 | 100661 | F | A | T | P | P | 372 | 384 |
| 2 | 12795684 | 100665 | F | C | T | NA | NA | 362 | 322 |
| 2 | 17808467 | 100665 | F | C | T | M | M | 362 | 322 |
| 2 | 54269316 | 100654 | F | G | T | M | M | 484 | 531 |
| 2 | 72750668 | 100665 | F | C | A | M | M | 362 | 322 |
| 2 | 74502317 | 100671 | F | A | C | P | P | 278 | 209 |
| 2 | 94627263 | 100657 | F | C | A | NA | M | 393 | 252 |
| 2 | 97253378 | 100664 | F | G | C | NA | NA | 237 | 197 |
| 2 | 105494061 | 100659 | F | G | A | P | P | 168 | 180 |
| 2 | 105754370 | 100646 | M | A | G | P | P | 286 | 167 |
| 2 | 124188222 | 100654 | F | A | T | P | P | 484 | 531 |
| 2 | 129040054 | 100657 | F | C | T | M | M | 393 | 252 |
| 2 | 135699807 | 100666 | F | G | A | P | P | 472 | 432 |
| 2 | 148074680 | 100646 | M | G | A | NA | M | 286 | 167 |
| 2 | 158818677 | 100654 | F | C | T | P | P | 484 | 531 |
| 2 | 167453781 | 100665 | F | C | T | P | P | 362 | 322 |
| 2 | 168940285 | 100648 | F | G | C | NA | NA | 190 | 167 |
| 2 | 171421375 | 100666 | F | A | C | M | M | 472 | 432 |

| | | | | | | | | | |
|---|---|---|---|---|---|---|---|---|---|
| 2 | 172439328 | 100664 | F | G | A | NA | P | 237 | 197 |
| 2 | 172439344 | 100664 | F | C | T | NA | P | 237 | 197 |
| 2 | 177146211 | 100653 | F | G | A | NA | NA | 398 | 445 |
| 2 | 203995431 | 100650 | F | A | G | NA | NA | 286 | 167 |
| 2 | 208497949 | 100649 | F | G | T | M | M | 270 | 247 |
| 2 | 245910010 | 100673 | F | C | A | P | P | 504 | 435 |
| 2 | 258495109 | 100666 | F | A | C | NA | NA | 472 | 432 |
| 2 | 260040592 | 100649 | F | A | G | P | P | 270 | 247 |
| 2 | 278101985 | 100660 | F | G | A | P | P | 293 | 305 |
| 2 | 296045008 | 100660 | F | C | T | P | P | 293 | 305 |
| 2 | 310844667 | 100662 | M | G | A | P | P | 393 | 252 |
| 2 | 379707031 | 100676 | F | G | T | NA | M | 198 | 293 |
| 2 | 429217257 | 100659 | F | G | A | M | M | 168 | 180 |
| 2 | 440862051 | 100659 | F | C | T | M | M | 168 | 180 |
| 2 | 446232330 | 100671 | F | A | G | NA | NA | 278 | 209 |
| 2 | 467627562 | 100672 | F | T | G | NA | NA | 390 | 321 |
| 2 | 487544564 | 100677 | F | G | A | NA | P | 323 | 418 |
| 2 | 493637682 | 100649 | F | C | A | M | M | 270 | 247 |
| 2 | 493637683 | 100649 | F | C | A | NA | M | 270 | 247 |
| 2 | 508197370 | 100677 | F | C | T | NA | NA | 323 | 418 |
| 2 | 528840436 | 100677 | F | C | A | NA | NA | 323 | 418 |
| 2 | 537087884 | 100648 | F | G | A | P | P | 190 | 167 |
| 2 | 541094310 | 100653 | F | G | A | P | P | 398 | 445 |
| 3 | 23156328 | 100673 | F | G | A | NA | NA | 504 | 435 |
| 3 | 30885082 | 100659 | F | C | T | M | M | 168 | 180 |
| 3 | 39599364 | 100669 | M | G | T | P | P | 328 | 146 |
| 3 | 40156225 | 100671 | F | G | A | P | P | 278 | 209 |
| 3 | 48650293 | 100650 | F | C | T | P | P | 286 | 167 |
| 3 | 87755540 | 100654 | F | T | C | P | P | 484 | 531 |
| 3 | 122866752 | 100666 | F | T | G | P | P | 472 | 432 |
| 3 | 163415496 | 100654 | F | G | A | P | P | 484 | 531 |
| 3 | 172633607 | 100664 | F | C | T | P | P | 237 | 197 |
| 3 | 180586861 | 100677 | F | T | C | NA | NA | 323 | 418 |
| 3 | 187355777 | 100677 | F | G | A | NA | NA | 323 | 418 |
| 3 | 198304210 | 100662 | M | T | C | P | NA | 393 | 252 |
| 3 | 241427634 | 100660 | F | A | T | NA | NA | 293 | 305 |
| 3 | 263894792 | 100652 | F | T | A | P | P | 298 | 345 |
| 3 | 275188809 | 100676 | F | G | A | M | M | 198 | 293 |
| 3 | 300278670 | 100672 | F | C | T | P | P | 390 | 321 |
| 3 | 308235691 | 100665 | F | T | C | P | P | 362 | 322 |
| 3 | 308235691 | 100666 | F | T | C | P | P | 472 | 432 |
| 3 | 330361550 | 100648 | F | C | A | NA | NA | 190 | 167 |
| 3 | 382053763 | 100649 | F | T | C | P | P | 270 | 247 |

| | | | | | | | | | |
|---|---|---|---|---|---|---|---|---|---|
| 3 | 389099082 | 100677 | F | G | A | P | P | 323 | 418 |
| 3 | 395246810 | 100653 | F | C | T | NA | NA | 398 | 445 |
| 3 | 412652068 | 100662 | M | A | T | NA | P | 393 | 252 |
| 3 | 480976412 | 100676 | F | G | A | P | P | 198 | 293 |
| 3 | 510989645 | 100669 | M | A | G | P | P | 328 | 146 |
| 3 | 521274310 | 100677 | F | A | C | M | M | 323 | 418 |
| 3 | 526036038 | 100646 | M | C | T | NA | M | 286 | 167 |
| 3 | 527559884 | 100660 | F | C | T | M | M | 293 | 305 |
| 4 | 31411469 | 100652 | F | C | A | M | M | 298 | 345 |
| 4 | 31411480 | 100652 | F | T | C | M | M | 298 | 345 |
| 4 | 46822034 | 100661 | F | C | T | NA | NA | 372 | 384 |
| 4 | 47146951 | 100672 | F | G | T | NA | NA | 390 | 321 |
| 4 | 57436414 | 100665 | F | C | G | M | M | 362 | 322 |
| 4 | 69376526 | 100660 | F | T | C | P | P | 293 | 305 |
| 4 | 73071609 | 100665 | F | A | G | M | M | 362 | 322 |
| 4 | 124495814 | 100654 | F | A | G | NA | P | 484 | 531 |
| 4 | 131590359 | 100676 | F | C | T | P | P | 198 | 293 |
| 4 | 141508971 | 100657 | F | G | T | P | P | 393 | 252 |
| 4 | 142336814 | 100677 | F | T | A | P | P | 323 | 418 |
| 4 | 151738120 | 100677 | F | A | G | P | P | 323 | 418 |
| 4 | 197376155 | 100676 | F | T | C | NA | M | 198 | 293 |
| 4 | 204798979 | 100648 | F | G | A | P | P | 190 | 167 |
| 4 | 212683994 | 100674 | F | G | A | P | P | NA | NA |
| 4 | 229466586 | 100660 | F | T | C | NA | NA | 293 | 305 |
| 4 | 234865629 | 100661 | F | T | C | P | P | 372 | 384 |
| 4 | 257499075 | 100652 | F | C | T | NA | NA | 298 | 345 |
| 4 | 283512603 | 100646 | M | G | A | M | M | 286 | 167 |
| 4 | 291317037 | 100653 | F | A | C | P | P | 398 | 445 |
| 4 | 298636685 | 100654 | F | G | A | NA | NA | 484 | 531 |
| 4 | 349954925 | 100676 | F | G | A | P | P | 198 | 293 |
| 4 | 352071991 | 100659 | F | G | C | NA | NA | 168 | 180 |
| 4 | 359530867 | 100659 | F | A | G | P | P | 168 | 180 |
| 4 | 359990184 | 100654 | F | G | A | P | P | 484 | 531 |
| 4 | 375228866 | 100674 | F | C | T | P | P | NA | NA |
| 4 | 385060918 | 100673 | F | G | T | P | P | 504 | 435 |
| 4 | 406588488 | 100674 | F | G | A | P | P | NA | NA |
| 4 | 408761655 | 100653 | F | A | G | NA | NA | 398 | 445 |
| 4 | 411613013 | 100666 | F | C | T | NA | P | 472 | 432 |
| 4 | 432139548 | 100648 | F | G | A | NA | NA | 190 | 167 |
| 4 | 433184699 | 100677 | F | G | T | P | P | 323 | 418 |
| 5 | 50942 | 100654 | F | C | T | P | P | 484 | 531 |
| 5 | 27691415 | 100669 | M | A | C | P | P | 328 | 146 |
| 5 | 45402218 | 100650 | F | G | T | P | P | 286 | 167 |

| | | | | | | | | | |
|---|---|---|---|---|---|---|---|---|---|
| 5 | 54069461 | 100652 | F | G | A | NA | NA | 298 | 345 |
| 5 | 66059896 | 100664 | F | C | T | NA | NA | 237 | 197 |
| 5 | 66256172 | 100674 | F | A | C | P | P | NA | NA |
| 5 | 76640686 | 100677 | F | T | A | M | M | 323 | 418 |
| 5 | 76640687 | 100677 | F | C | T | NA | M | 323 | 418 |
| 5 | 85086709 | 100666 | F | T | C | M | M | 472 | 432 |
| 5 | 106038059 | 100660 | F | T | A | NA | NA | 293 | 305 |
| 5 | 114632632 | 100664 | F | A | C | NA | NA | 237 | 197 |
| 5 | 138475884 | 100674 | F | C | T | M | M | NA | NA |
| 5 | 142038799 | 100673 | F | G | T | M | M | 504 | 435 |
| 5 | 146963478 | 100665 | F | C | A | M | M | 362 | 322 |
| 5 | 149037863 | 100676 | F | G | A | M | M | 198 | 293 |
| 5 | 182182265 | 100650 | F | G | A | M | M | 286 | 167 |
| 5 | 193699510 | 100669 | M | A | C | P | P | 328 | 146 |
| 5 | 198660922 | 100671 | F | T | G | P | P | 278 | 209 |
| 5 | 200251220 | 100665 | F | G | T | NA | NA | 362 | 322 |
| 5 | 200978502 | 100659 | F | T | G | P | P | 168 | 180 |
| 5 | 208339338 | 100676 | F | G | A | P | P | 198 | 293 |
| 5 | 214110070 | 100660 | F | T | C | P | P | 293 | 305 |
| 5 | 253139723 | 100659 | F | C | T | P | P | 168 | 180 |
| 5 | 266882209 | 100653 | F | T | A | NA | NA | 398 | 445 |
| 5 | 266882209 | 100654 | F | T | A | NA | NA | 484 | 531 |
| 5 | 292373558 | 100662 | M | C | A | NA | P | 393 | 252 |
| 6 | 904321 | 100672 | F | C | G | NA | NA | 390 | 321 |
| 6 | 1741611 | 100666 | F | C | T | NA | NA | 472 | 432 |
| 6 | 10892497 | 100657 | F | T | G | P | P | 393 | 252 |
| 6 | 16603265 | 100671 | F | G | A | NA | NA | 278 | 209 |
| 6 | 23926088 | 100649 | F | A | T | P | P | 270 | 247 |
| 6 | 37088581 | 100654 | F | T | A | P | P | 484 | 531 |
| 6 | 55004062 | 100676 | F | G | A | P | P | 198 | 293 |
| 6 | 68851283 | 100671 | F | G | A | P | P | 278 | 209 |
| 6 | 120046271 | 100665 | F | C | T | NA | NA | 362 | 322 |
| 6 | 128971367 | 100659 | F | A | G | M | M | 168 | 180 |
| 6 | 128971368 | 100659 | F | C | A | NA | M | 168 | 180 |
| 6 | 169392874 | 100657 | F | A | C | M | M | 393 | 252 |
| 6 | 189833370 | 100672 | F | C | T | P | P | 390 | 321 |
| 6 | 195480675 | 100677 | F | A | G | P | P | 323 | 418 |
| 6 | 207424315 | 100653 | F | G | A | M | M | 398 | 445 |
| 6 | 237109016 | 100666 | F | A | C | NA | NA | 472 | 432 |
| 6 | 249648594 | 100646 | M | C | T | P | P | 286 | 167 |
| 6 | 250446646 | 100676 | F | T | C | NA | M | 198 | 293 |
| 6 | 267924003 | 100648 | F | A | G | NA | P | 190 | 167 |
| 6 | 278479241 | 100665 | F | T | A | NA | NA | 362 | 322 |

| | | | | | | | | | |
|---|---|---|---|---|---|---|---|---|---|
| 7 | 23250608 | 100661 | F | G | T | M | M | 372 | 384 |
| 7 | 67824981 | 100654 | F | T | C | M | M | 484 | 531 |
| 7 | 115570795 | 100672 | F | C | T | NA | NA | 390 | 321 |
| 7 | 122505580 | 100653 | F | T | C | M | M | 398 | 445 |
| 7 | 124705728 | 100653 | F | T | C | M | M | 398 | 445 |
| 7 | 152612830 | 100666 | F | C | A | P | P | 472 | 432 |
| 7 | 173554604 | 100664 | F | C | G | P | P | 237 | 197 |
| 7 | 190291332 | 100671 | F | C | T | P | P | 278 | 209 |
| 7 | 213699398 | 100649 | F | G | A | P | P | 270 | 247 |
| 7 | 222175329 | 100660 | F | C | T | P | P | 293 | 305 |
| 7 | 236485705 | 100659 | F | G | C | M | M | 168 | 180 |
| 7 | 240611322 | 100669 | M | T | A | P | P | 328 | 146 |
| 7 | 244711985 | 100649 | F | C | T | P | P | 270 | 247 |
| 7 | 246290255 | 100654 | F | C | G | NA | NA | 484 | 531 |
| 8 | 12490958 | 100677 | F | G | A | P | P | 323 | 418 |
| 8 | 14174448 | 100677 | F | A | G | P | P | 323 | 418 |
| 8 | 17767006 | 100677 | F | A | G | NA | NA | 323 | 418 |
| 8 | 42462187 | 100666 | F | G | T | P | P | 472 | 432 |
| 8 | 48925131 | 100661 | F | C | T | NA | NA | 372 | 384 |
| 8 | 53644751 | 100671 | F | A | G | P | P | 278 | 209 |
| 8 | 68176410 | 100660 | F | A | G | P | P | 293 | 305 |
| 8 | 92644000 | 100676 | F | A | G | NA | NA | 198 | 293 |
| 8 | 93628298 | 100659 | F | C | T | NA | P | 168 | 180 |
| 8 | 110071417 | 100657 | F | G | A | M | M | 393 | 252 |
| 8 | 135163984 | 100676 | F | T | G | NA | NA | 198 | 293 |
| 8 | 140693709 | 100648 | F | G | A | P | P | 190 | 167 |
| 8 | 191490323 | 100671 | F | C | T | NA | NA | 278 | 209 |
| 8 | 201562122 | 100669 | M | T | C | M | M | 328 | 146 |
| 8 | 201562122 | 100674 | F | T | C | M | M | NA | NA |
| 8 | 213832244 | 100654 | F | G | T | NA | M | 484 | 531 |
| 8 | 213832267 | 100654 | F | C | T | M | M | 484 | 531 |
| 8 | 213832268 | 100654 | F | A | G | NA | M | 484 | 531 |
| 8 | 229502907 | 100660 | F | G | A | M | M | 293 | 305 |
| 8 | 234830176 | 100648 | F | G | T | P | P | 190 | 167 |
| 8 | 236353340 | 100653 | F | T | C | P | P | 398 | 445 |
| 8 | 270801073 | 100659 | F | C | T | P | P | 168 | 180 |
