## Supplementary material for "Low mutation rate but high male-bias in the germline of a short-lived opossum": Figure S1

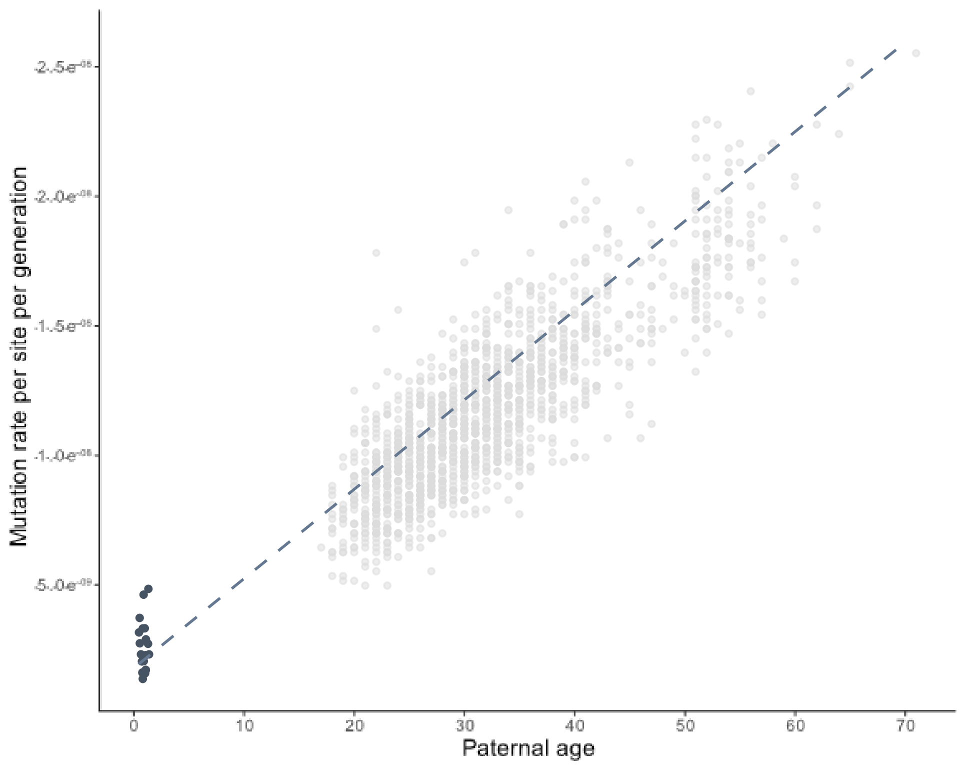


**Figure S1.** Low mutation rate due to early reproductive age. The predicted mutation rate in opossums (blue dots, one per trio) is derived from regression estimates based on human mutation data (gray dots) and adjusted for parental age and callable genome size. The dashed line represents the regression fit from human trios at different paternal ages.
