## Supplementary material for "Low mutation rate but high male-bias in the germline of a short-lived opossum": Figure S2

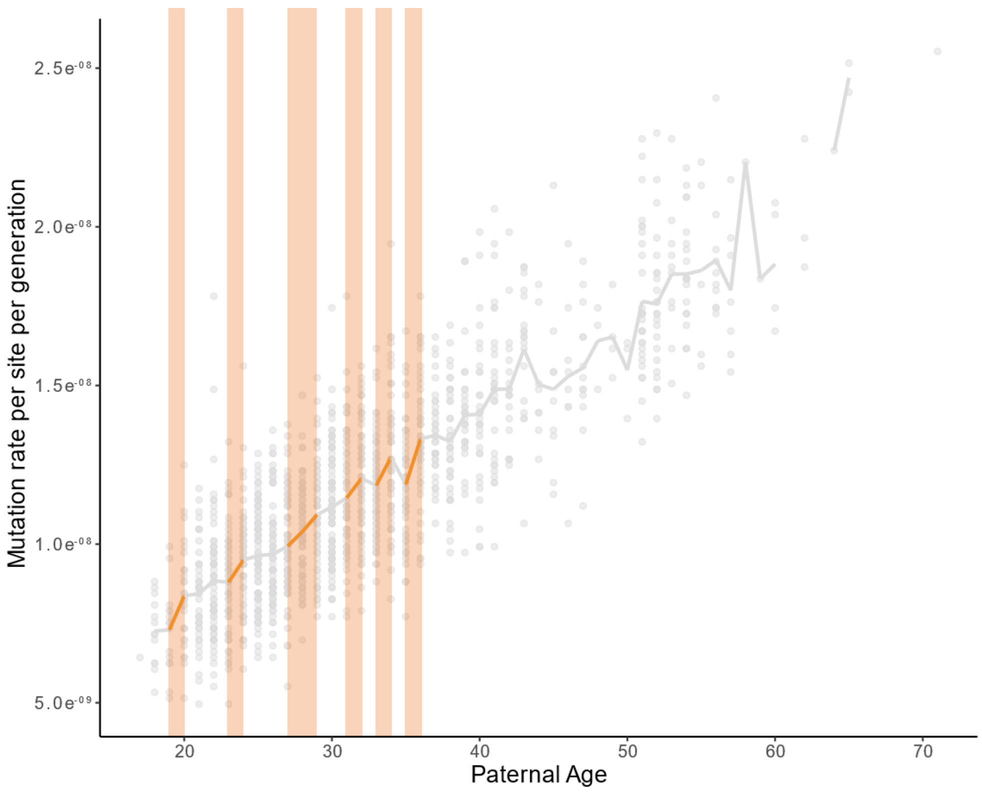


**Figure S2.** Paternal age effect in human mutation rates using a restricted age range. Each gray dot represents the mutation rate observed at a specific paternal age. Consecutive paternal age pairs, separated by only one year, were analyzed to assess whether a significant age effect exists. Only 7 of 44 consecutive age pairs (highlighted in orange) show a significant paternal age effect, suggesting that the absence of a paternal age effect in opossums may be due to the narrow age range among parents.
