## Supplementary material for "Low mutation rate but high male-bias in the germline of a short-lived opossum": Figure S3

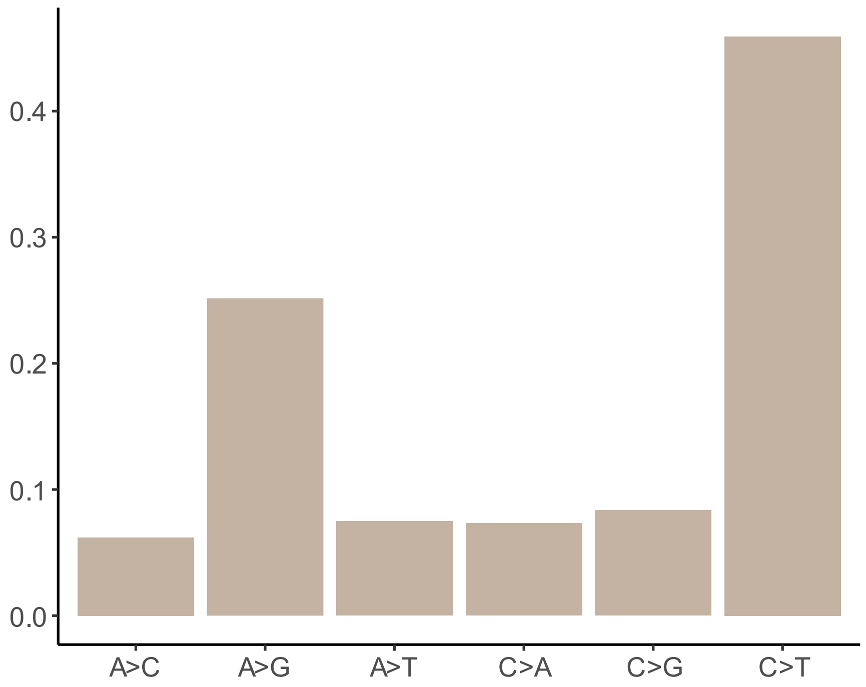


**Figure S3.** Predicted mutation spectrum in opossums. The predicted mutation spectrum in opossums illustrates the expected mutation counts for each mutation class. Each class is modeled as a function of maternal and paternal age, accounting for typical Poisson-distributed mutation rates.
