## Supplementary material for "Low mutation rate but high male-bias in the germline of a short-lived opossum": Figure S4

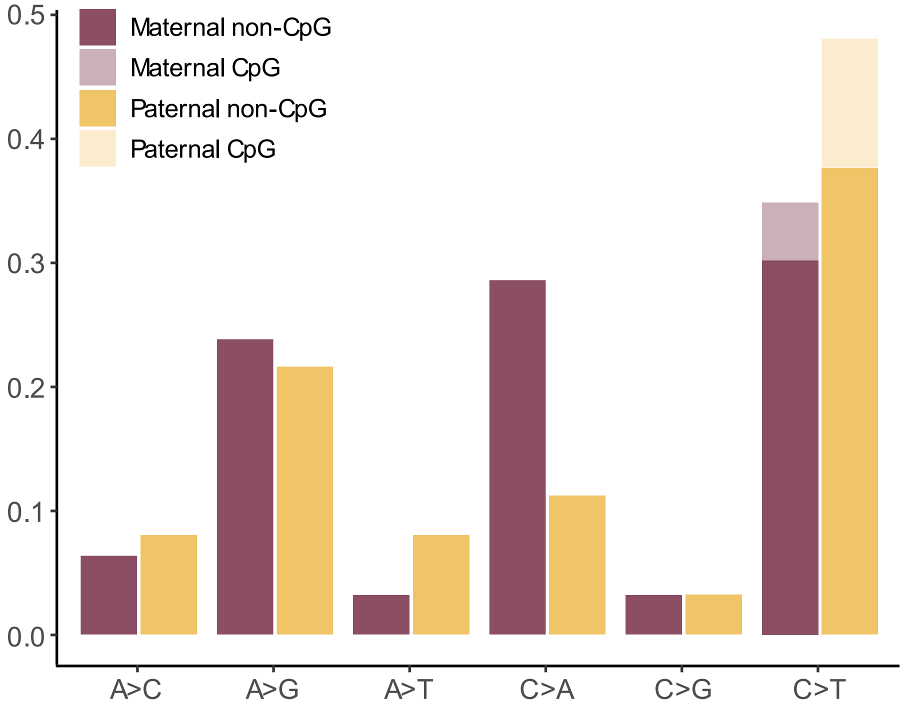


**Figure S4.** Sex-Specific Mutation Spectrum in Opossums. The sex-specific mutation spectrum in opossums was analyzed using all 188 phased mutations, which accounted for 76.4% of the total 246 mutations, to evaluate variation in the types of mutations inherited from mothers versus fathers.
